# Cultivation protocols for the rhodophytes, *Devaleraea mollis* and *Palmaria hecatensis* from Alaska

**DOI:** 10.1101/2024.10.20.619297

**Authors:** Muriel C. Dittrich, Lindsay Meyer, Michael Stekoll, Amanda Kelley, Schery Umanzor

## Abstract

Species diversification is crucial for the long-term viability, competitiveness, and sustainability of the seaweed farming industry in the United States. This study investigated the effects that temperature (4, 8 and 12°C), photoperiod (8L:16D, 12L:12D and 16L:8D), and irradiance (20, 40, 100 ± 10 µmol photons m^−2^ s^−1^) had on the specific growth rate (SGR) of *Devaleraea mollis* and *Palmaria hecatensis* from Alaska. Outputs were used to adjust indoor cultivation protocols for *D. mollis* and develop the first protocols for *P. hecatensis*. This study also explored the use of two low-cost commercial nutrient products (F/2 and Jack’s Special 25-5-15) as alternatives to the recommended nutrient medium, von Stosch Enrichment. Assessments determined whether their use resulted in similar SGRs without compromising biomass quality or raising production costs. Results showed that both species responded differently to each factor, indicating distinct ecological and physiological adaptations. *D. mollis* exhibited higher growth rates in warmer temperatures and responded to higher irradiance levels with spore release but showed no clear preference between neutral and long-day photoperiods. In contrast, *P. hecatensis* demonstrated higher growth rates in cooler environments with a long-day photoperiod promoting the most growth without spore release. Nutrient supplementation revealed that growth in *D. mollis* was affected by nutrient formulation, while *P. hecatensis* showed no significant growth variation. These findings underscore the importance of species and geographic-specific protocols for seaweed farming. Further research is needed to optimize the potential cultivation protocols provided here. Cultivation protocols would also benefit from exploring the ecological and physiological nuances of both species.

## Introduction

The global seaweed industry, valued at approximately USD 17 billion (FAO 2024), spans diverse species and uses. Around 95% of this value stems from seaweed cultivated for human consumption, underscoring its essential role in global diets (FAO 2024). Hydrocolloid recovery represents the second largest sector of the market, while animal feed additives and fertilizers make up the rest (FAO 2016; Holdt and Kraan 2011). Additionally, the industry is pursuing new opportunities in high-value sectors such as cosmetics, pharmaceuticals, biomedical products, and nutraceuticals (Edwards and Dring 2011;Winberg et al. 2011; FAO 2018). The increasing demand for seaweed biomass is fueled by the appeal of sustainable food options and comprehensive health benefits (Wu et al. 2023). Projections suggest that the seaweed market could grow by an additional USD 11.8 billion by 2030, reflecting substantial potential within this expanding industry (World Bank 2023). Specifically, farming of red seaweed holds significant commercial interest as several species are rich in hydrocolloids (e.g., agar and carrageenan), secondary metabolites, and protein (Hafting et al. 2012; FAO 2014; FAO 2016). Red seaweeds are also rich in phycobiliproteins, specifically phycoerythrin, and phycocyanin, which are pigment-protein complexes (Freitas et al. 2021). These pigments are widely utilized in scientific, medical, and commercial applications due to their bright fluorescence and nutraceutical properties (Nowruzi and Hashemi 2024). Phycoerythrin is commonly used in flow cytometry, fluorescence microscopy, and immunoassays such as ELISAs for enhanced sensitivity and accuracy in antigen or antibody detection (Smith et al. 2019; Chen et al. 2023). Phycocyanin is used as a fluorescent marker in various bioassays and imaging techniques for detecting and analyzing biological molecules and cells (Gadberry et al. 2018; Freitas et al. 2021). Beyond biotechnology, phycocyanin is used in the food and cosmetic industries as a natural colorant and antioxidant, offering putative anti-inflammatory and immune-boosting effects (Fernandes et al. 2023). These versatile and high-end applications underscore the need to increase and improve access to red seaweed biomass to satisfy demands for research, clinical diagnostics, and commercial products.

To date, red seaweeds account for approximately 53% of the total cultivated seaweed biomass globally (FAO 2024). Most of this biomass is produced in Asian countries, although efforts are underway to increase production in Africa, America, and Europe (FAO 2024). In the United States, experimental farming of red seaweeds started in the early 1990s (Mumford 1990; Yarish et al. 1998), with successful long-term and small-scale commercial operations beginning approximately 15 years later (Kim et al. 2019). Among the species of interest, *Gracilaria tikvahiae* and *Palmaria palmata* on the East Coast, and *Devaleraea mollis* (formerly known as *Palmaria mollis*) and *Palmaria hecatensis* on the West Coast, stand out as top candidates for human consumption. In particular, *P. palmata* and *D. mollis*, are of interest due to their high protein content, ranging from 8-35% of dry weight (Buckley and Stekoll 2006).

*D. mollis* has been successfully grown in outdoor land-based systems in Oregon and California. Extensive work has examined the relationship between biomass production and environmental parameters (Rosen et al. 2000; Demetropoulos and Langdon 2004a; Demetropoulos and Langdon 2004b). Cultivation of the species has gained traction, given it is often described as the “bacon seaweed” and is currently used as a vegan alternative to animal-based bacon (e.g., Umaro Foods). No cultivation efforts have been described for *P. hecatensis*. Significant financial investment in the last decade has enabled Alaska to emerge as a seaweed farming hub. The state is now second in production, following Maine (Robidoux & Good 2023). Alaska aims to develop a $100 million mariculture industry by 2041 (Alaska Mariculture Task Force 2018). The focus has been primarily on kelp and oyster farming at sea. However, on-land cultivation of red seaweed could offer new market opportunities and help Alaska achieve its industry goal within the next decade.

In Alaska, *D. mollis* and *P. hecatensis* hold cultural significance among Tlingit communities. These species are essential to the diets of coastal Indigenous communities and also serve as valuable trade goods (Garza et al. 1989; Johnson et al. 2019). Wild populations of *D. mollis* can be found year-round in certain locations such as Cook Inlet, Southcentral Alaska, where it colonizes mariculture gear (W. Bates, pers. comm.). This year-round availability differs from the seasonal life cycle usually described for the species, which spans from early spring to summer (Hawkes 1985). There is limited information related to the ecology of *P. hecatensis*. However, studies in British Columbia, Canada, describe the potential for perennation of older blades (Hawkes 1985), though this has yet to be confirmed for Alaska and other regions. While *D. mollis* has a geographical distribution that expands from Southern California to the Aleutian Islands, Alaska, *P. hecatensis* has a more limited presence with patches originally described for Southeast Alaska, British Columbia, Canada, and the eastern coast of Russia (Skriptsova et al. 2023).

Early attempts to grow Alaskan ecotypes of *D. mollis* following cultivation protocols used in California and Oregon (Oregon Seaweed & Monterey Bay Seaweed) were unsuccessful. The failures were attributed to unsuitable temperature and light conditions. Natural coastal conditions in Alaska differ significantly from those in California and Oregon, where the current protocols were developed. For example, Alaska experiences sharper changes in seawater temperature and photoperiod (ratio of light-to-dark hours in a 24-hr. period) than any other state in lower latitudes. Therefore, cultivation protocols adjusted to conditions naturally experienced in Alaska must be developed as a first step to start the commercial on-land and indoor cultivation of *D. mollis*. Conversely, *P. hecatensis* is a potential novel mariculture species. It shares similarities with *D. mollis* in its life cycle, habitable range, traditional use in local diets, and potential as a feed source in abalone farming, meeting protein needs and supplying biomass for high-value markets.

Unlike open-sea cultivation, indoor systems allow precise control over environmental factors, including light, temperature, salinity, pH, water turbulence, and nutrient concentrations (Redmond et al. 2014). This controlled environment supports customized biomass production, ensuring biosafety and meeting high-quality standards tailored to market demands (Mendoza et al. 2018). In this study, we measured the effects of temperature, photoperiod, and irradiance on the specific growth rate (SGR) of *D. mollis* and *P. hecatensis* from Alaska. Outputs were used to adjust indoor cultivation protocols for *D. mollis* and develop the first protocols for *Palmaria hecatensis*. After establishing suitable growth conditions, the study compared the effect of three nutrient sources on growth for both species. Assessments focused on determining whether their use resulted in similar SGRs without compromising biomass quality or raising production costs. Our broader objective was to diversify Alaska’s mariculture portfolio by expanding species options for sustainable indoor cultivation.

## Materials and Methods

### Collection and acclimation of stocking biomass

Experimental *Devaleraea mollis* and *Palmaria hecatensis* were collected from Middle Bay, Kalsin Bay, and Pasagshak Bay in Kodiak Island, Alaska, between March and April 2022 and 2023 when biomass abounds (Fig. 1). Approximately 25 kg of healthy thalli were detached from the intertidal zone at low tide (−0.5 m MLLW). Thalli with proliferations, minimal herbivory, and no signs of decay or bleaching were selected (Fig. 2). Biomass was wrapped in seawater-damped paper towels, packed into sealed plastic tubs with gel ice packs, and transported in coolers to the College of Fisheries and Ocean Sciences, University of Alaska Fairbanks in Juneau. Transit time did not exceed 24 hours.

**Fig. 1.**
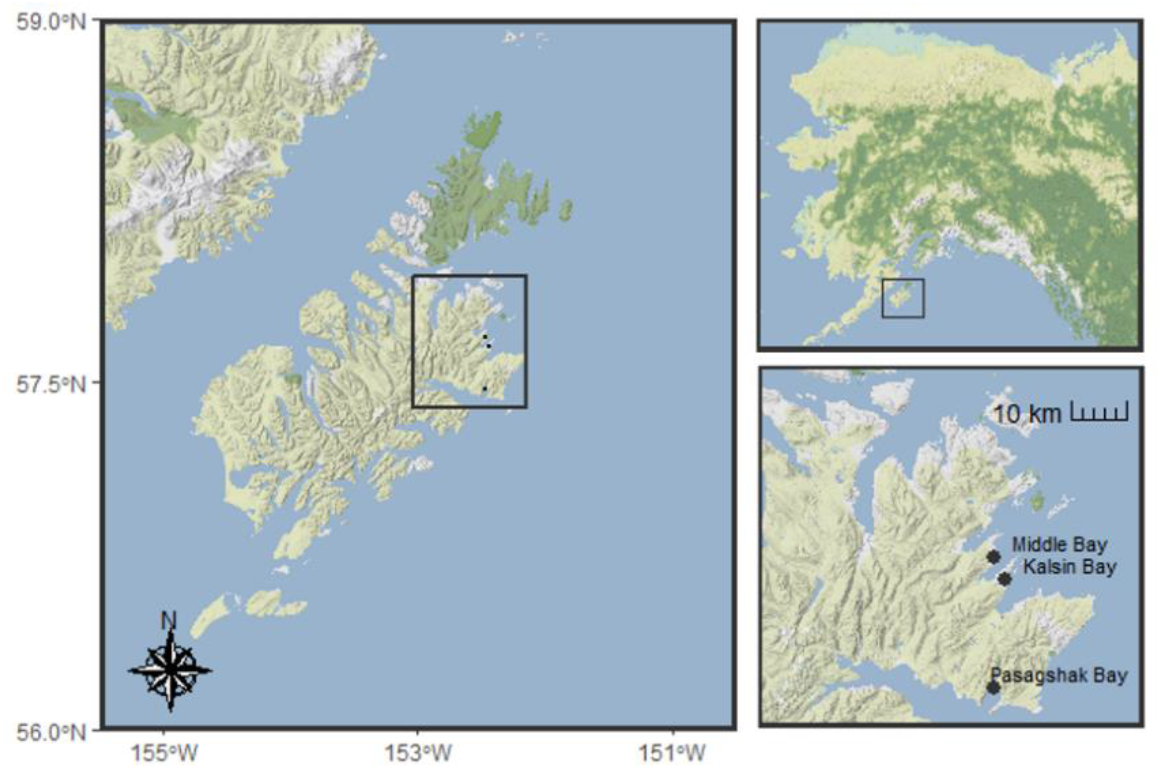
Collection sites (Middle, Kalsin, and Pasagshak Bays) on the east side of Kodiak Island in Southcentral Alaska.

**Fig. 2.**
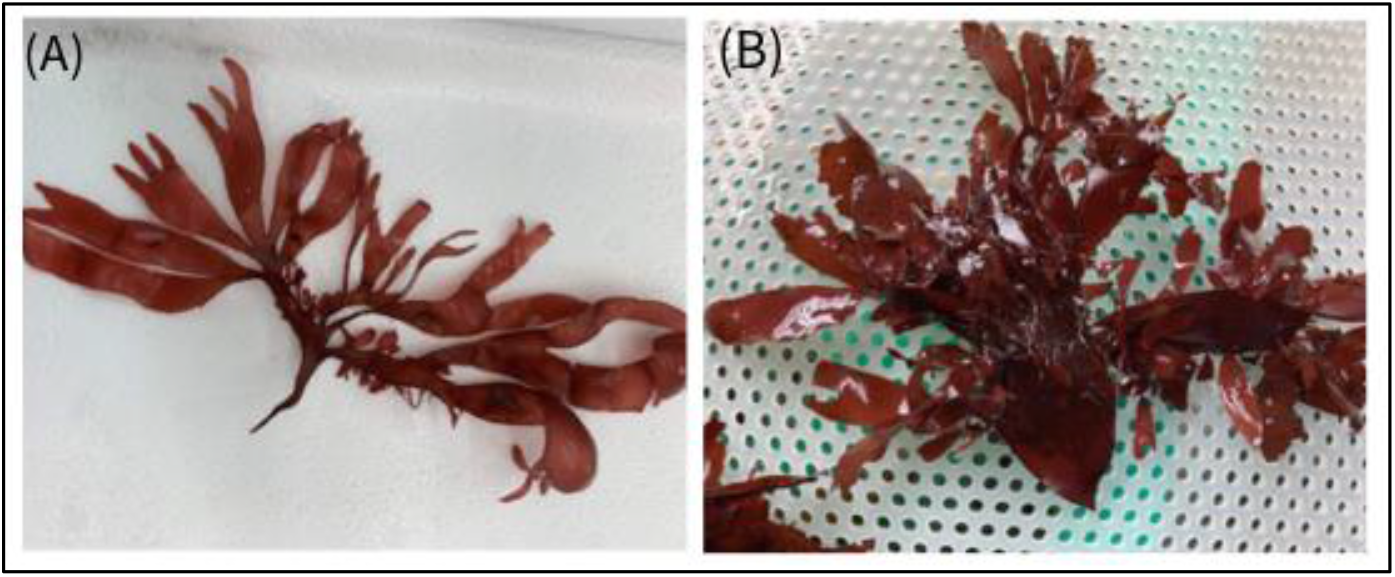
(A) *Devaleraea mollis* and (B) *Palmaria hecatensis* showing multiple proliferations on the edge and middle section of their thalli.

Upon arrival, thalli were placed in independent 1000L recirculating tanks (30 ± 1 ppt salinity, 4 ± 0.5°C, 50 ± 10 µmol photons m^−2^ s^−1^, 12L:12D) for 14 days to acclimate. Tank conditions were set similar to those at the collection sites. Tanks were cleaned and refilled with filtered (1 µm) and UV-treated seawater every seven days. A portion of the biomass was used for experiments, while the rest was maintained under the same conditions as stock for future experiments.

### Preparation of experimental samples

After the 14-day acclimation, 90 proliferations per species were detached from at least ten parental thalli. Proliferations were rinsed with sterile seawater (SSW), rubbed with a Kimwipe to remove surface contaminants, and grouped in sets of three (0.01001 g ± 0.006 g). Each group was placed into 250 mL Erlenmeyer flasks filled with SSW (30 ± 1 ppt) enriched with F/2 and 2mL/L germanium dioxide (250 mg L^-1^ stock) to control diatoms (Lewin 1965). Flasks were placed in Conviron Growth Chambers (Conviron GE1000) set to 4°C, 40 ± 10 µmol photons m^−2^ s^−1^, 12L:12D, with experimental conditions adjusted as needed. Flasks were aerated throughout the experiment. See Figure 3 for a diagram of the experimental workflow.

**Fig. 3.**
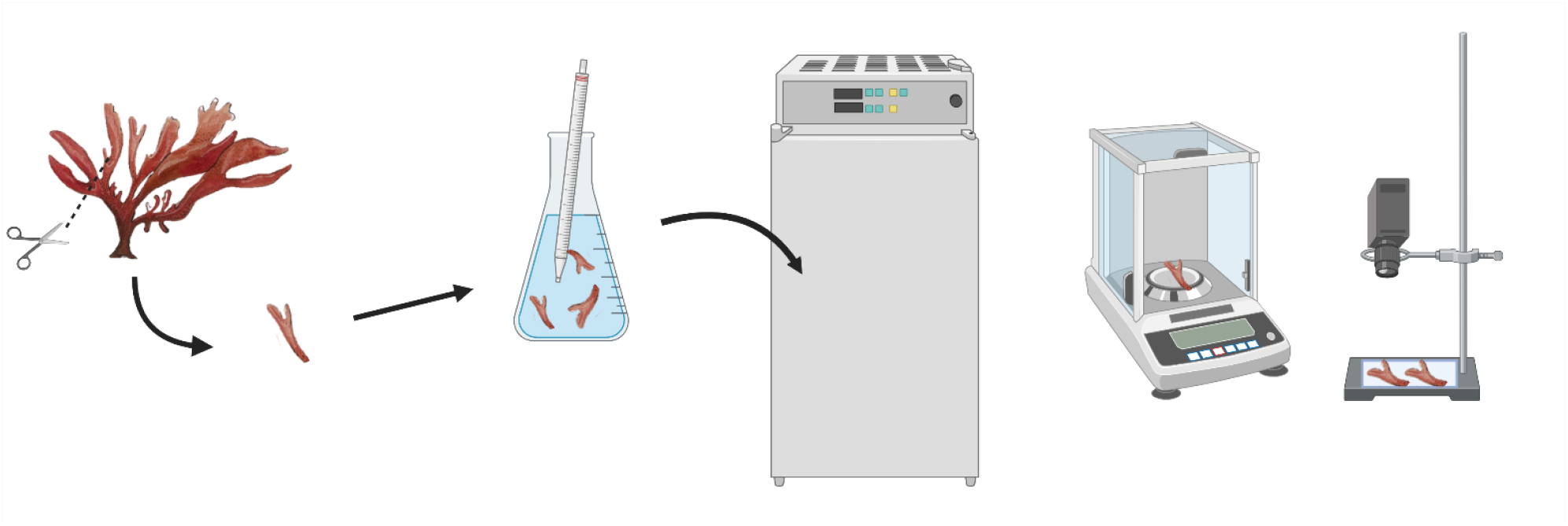
Experimental workflow including selection of proliferations (samples), allocation into 250mL Erlenmeyer flasks (n=5), exposures to treatments inside growth chambers, weighing, and photographing. Created in BioRender.

### Experimental Design

The experiments exposed *D. mollis* (n = 5) and *P. hecatensis* (n = 5) to three temperatures (4, 8 and 12°C), photoperiods (16L:8D, 12L:12D, 8L:16D), and irradiances (20, 40, 100 ± 10 µmol photons m^−2^ s^−1^) to identify grow conditions in indoor settings. Growth was measured using wet weight as the metric.

Temperature and photoperiod were tested on all possible combinations. However, given the availability of only three incubators, culture conditions were adjusted such that only one experimental factor was manipulated at a time, using a stepwise approach rather than a full factorial setup. Irradiance was tested using the temperature and photoperiod, rendering the most thalli growth, with no tissue paleness or signs of fertility. Each experiment lasted 28 consecutive days. Fresh proliferations were used for each factor tested. Experiments were conducted in 2022 and 2023.

### Experiments

Three growth chambers were used throughout the experiments. Initial trials determined upper-temperature limits, showing thalli bleached at 15°C but continued growing at 12°C, though pale. Thus, temperatures were set at 4, 8, and 12°C, matching water temperatures in Southeast and Southcentral Alaska. Samples were conditioned to their experimental temperatures by increasing 2°C every 24 hours, with photoperiod and irradiance held constant (12L:12D, 40 ± 10 µmol photons m^2^ s^−1^). Cultures were enriched with F/2 every seven days to avoid nutrient limitation. Growth data were recorded every seven days from pat-dried biomass placed on an analytical balance (Metter Toledo ME204E). Once weighed, biomass was photographed to record potential color changes that could indicate nutrient limitation or light stress (Sahoo and Yarish 1996) and morphology or texture potentially indicating fertility or asexual reproduction. After that, biomass returned to their flask, where they continued to grow until the completion of the experimental cycle. The same data collection process was repeated for photoperiod and irradiance. Each parameter was assessed using fresh proliferations at the start of the experiments. Biomass was normalized to obtain SGRs, following the equation provided by Shin et al. (2020):

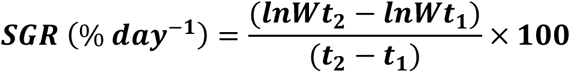

Photoperiod levels resembled natural light-to-dark ratios (8L:16D, 12L:12D, 16L:8D) during the seaweed natural growing period from late February to July. Each photoperiod was tested at each temperature following a stepwise approach, while irradiance remained at 40 ± 10 µmol photons m^2^ s^−1^. Cultures were enriched with F/2 every seven days to avoid nutrient limitation. Irradiance (20, 40, 100 ± 10 µmol photons m^−2^ s^−1^) were selected based on existing growing protocols for *D. mollis* (Demetropoulos and Langdon 2004a). Irradiance was gradually adjusted by changing photon fluence 10 µmol photons m^−2^ s^−1^ every 6 hours until the desired setups were reached. These adjustments were completed at a suitable temperature and photoperiod determined in the previous experiment (8°C and 16L:8D). Subsequent evaluations tested the effect of von Stosch Enrichment medium (VSE), Guillard’s f/2 medium (F/2), and Jack’s Special 25-5-15 (JS) on growth. These nutrient sources were selected based on accessibility, cost, and formulation, particularly focusing on the molarity of available nitrogen (Table 1). Culture conditions utilized a temperature of 8°C, 12L:12D photoperiod, and 100 ± 10 µmol photons m^2^ s^−1^ for both species.

**Table 1.** Chemical composition of Von Stosch enrichment, (VSE) F/2, and Jack’s Special (JS) used as experimental nutrient sources. Modified from Kim and Yarish 2014.

| Nutrient ($\mu\text{M}$) | Von Stosch Enrichment Medium (VSE) | Guillard's F/2 Medium (F/2) | Jack's Special (N-P-K, 25-5-15) (JS) |
| --- | --- | --- | --- |
| Nitrogen | 500<br>(100% $\text{NO}_3^-$) | 500<br>(100% $\text{NO}_3^-$) | 500<br>(57% $\text{NO}_3^-$ and 43% $\text{NH}_4^+$) |
| Phosphorus | 30 | 15 | 45 |
| Iron | 1 | 11.7 | 0.6 |
| Manganese | 10 | 0.9 | 0.3 |
| EDTA | 10 | 11.7 | - |
| Vitamins | Vitamin B<sub>1</sub>, Biotin, Vitamin B<sub>12</sub> | Vitamin B<sub>1</sub>, Biotin, Vitamin B<sub>12</sub> | - |
| Potassium | - | - | 107 |
| Boron | - | - | 0.6 |
| Copper | - | 0.04 | 0.05 |
| Molybdenum | - | 0.03 | 0.03 |
| Zinc | - | 0.05 | 0.004 |
| Magnesium | - | - | 1.15 |

A biomass threshold of 1 g L^-1^ was set to prevent carbon, nutrient, light, or flow limitations. Thalli were transferred from 250 mL flasks to 500 mL or 1 L flasks when this threshold was met (Redmond et al. 2014). Water changes, conducted immediately after weighing and photographing, involved replacing the flask with filtered, UV-treated, and autoclaved seawater pre-adjusted to the desired temperature. After establishing suitable cultivation protocols, the experimental biomass was gradually scaled up from 250 mL flasks to a 150 L cylindrical fiberglass tank.

### Data analysis

Differences in SGRs from days 0 to 28 across temperature, photoperiod, irradiance, and nutrient source treatments were analyzed using one-way ANOVAs. Before analysis, assumptions were validated. Normality was assessed with the Shapiro-Wilk test, and homogeneity of variances was checked using Bartlett’s test. Welch’s one-way ANOVA was applied for cases where assumptions were not met.

Significance was set at α = 0.05, and post-hoc comparisons included Tukey’s HSD, Games-Howell test, and pairwise t-tests with Bonferroni corrections to control for Type I error. Weekly SGR changes per factor and species were visualized using line graphs added to the Appendix section.

## Results

### Devaleraea mollis

The effect of temperature on SGRs for *D. mollis* varied with photoperiod (p < 0.05; Tables S1–S3; Fig. 4). Across all temperatures tested, SGRs were consistently higher at photoperiods of 16L:8D and 12L:12D compared to 8L:16D (Tukey’s HSD p < 0.001; Table S4). Under a 16L:8D photoperiod, SGRs averaged around 9.8% per day at 8 and 12°C (Tukey’s HSD, p < 0.05; Fig. 4, Table S2), while under a 12L:12D photoperiod, SGRs remained approximately 11% daily across temperatures. SGRs for the 12L:12D and 16L:8D photoperiods differed only at 4°C, where SGRs were significantly higher at 12L:12D (Tukey’s HSD, p < 0.001; Table S4).

**Fig. 1.**
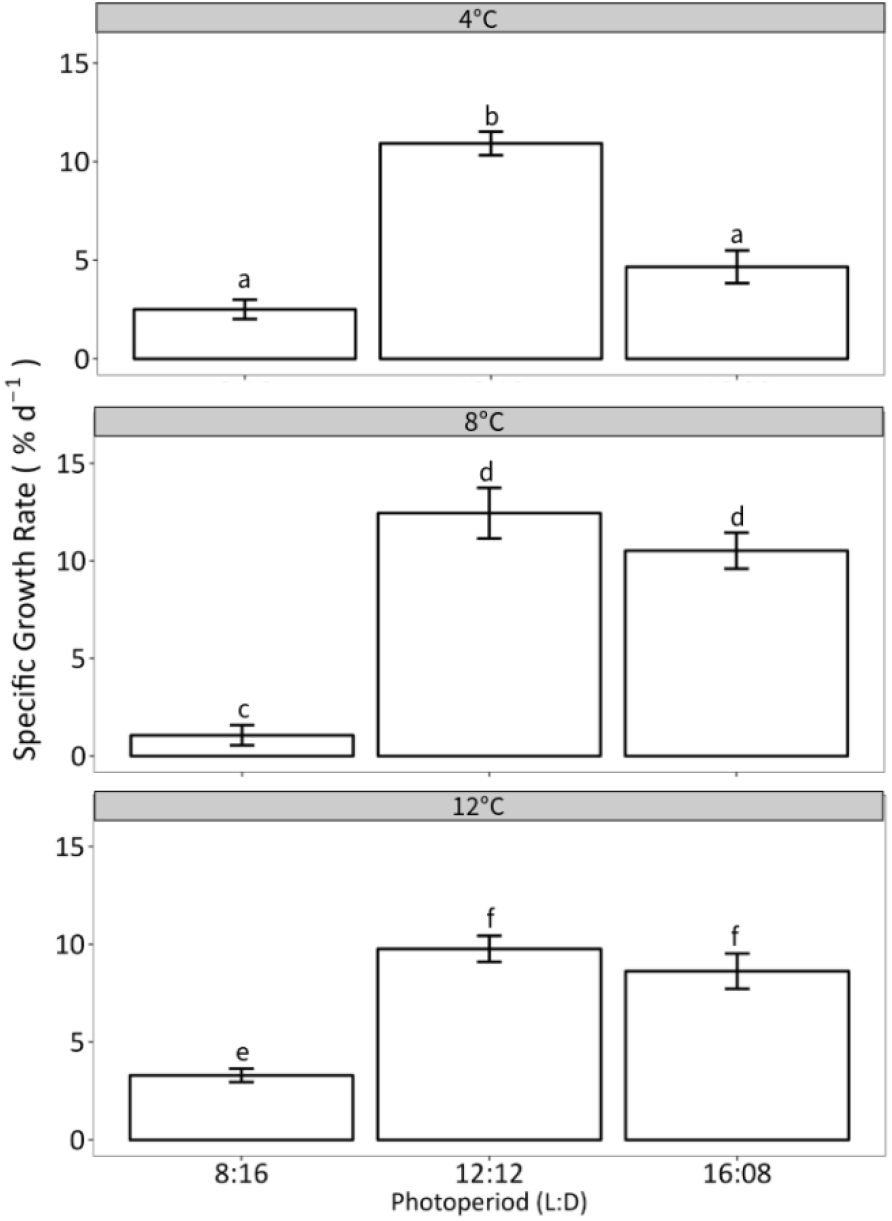
(A) Specific growth rate of *Devaleraea mollis* as a function of temperature (4, 8, and 12°C) and photoperiod (16L:08D, 12L:12D, and 8L:16D) grown with 40 µmol photons m^-2^ s^-1^ and F/2 as the nutrient source. Experiments were run following a stepwise approach. Data shows mean values ± SE, n = 5. Significance is denoted by letters.

At the 12L:12D photoperiod, SGRs remained above 9% per day across all temperatures. Similarly, SGRs at 16L:8D were above 8% daily at 8 and 12°C but dropped to around 5% at 4°C. In contrast, when thalli were grown under an 8L:16D photoperiod, SGRs did not exceed 4% regardless of temperature (Fig. 4 and Fig. S1).

Similar to temperature and photoperiod, SGRs varied significantly with irradiance levels when combined with 8°C and a 16L:8D photoperiod (p < 0.001; Fig. 5A; Table S5). SGRs were highest in samples exposed to an irradiance of 100 µmol photons m^−2^ s^−1^ (Tukey’s HSD p < 0.0001; Table S5), followed by 40 µmol photons m^−2^ s^−1^ and 20 µmol photons m^−2^ s^−1^, each of which differed significantly from one another (Tukey’s HSD p < 0.01; Table S5). Biomass exposed to 100 µmol photons m^−2^ s^−1^ displayed fertility signs by day 14. Fertility was indicated by changes in thalli color and texture, followed by spore release by day 21. This reproductive activity resulted in a 9% reduction in SGR that week (see Fig. S2).

**Fig. 2.**
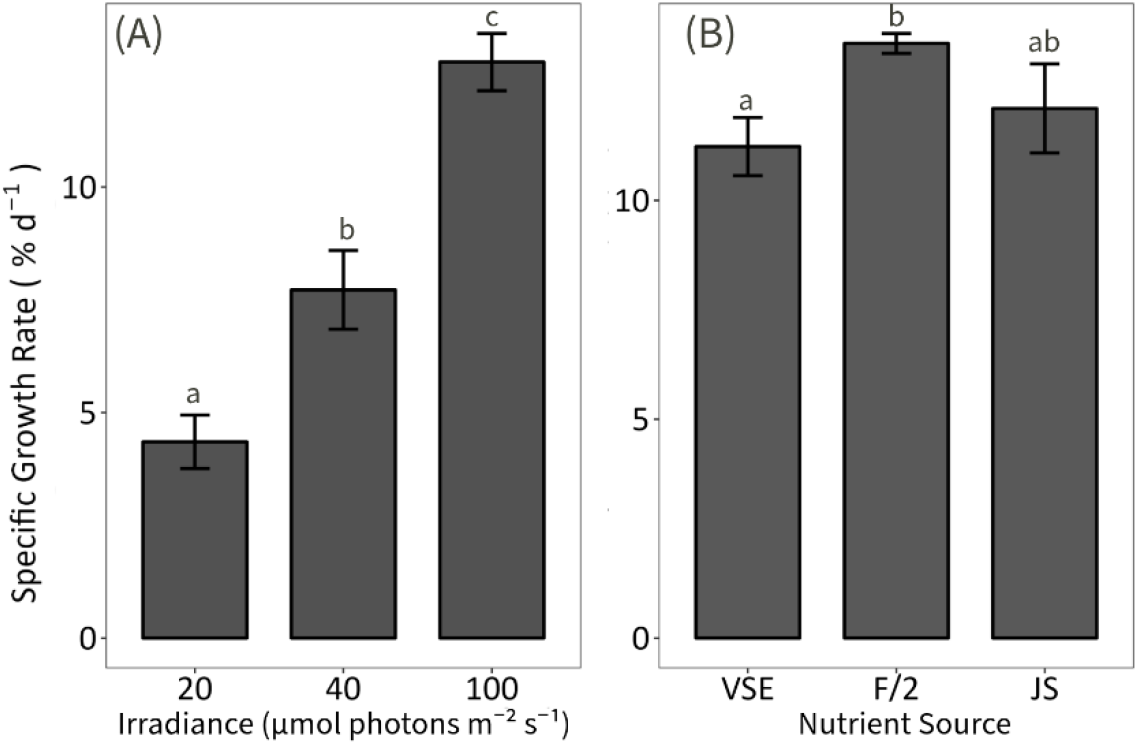
Specific growth rate of *Devaleraea mollis* as a function of (A) irradiance (20, 40, and 100 µmol photons m^-2^ s^-1^) with 16L:08D, 8 °C and F/2 as the nutrient source, n = 5. (B) SGR as a function of nutrient source, with 8 °C, 16L:8D, and 40 µmol photons m^-2^ s^-1^, n = 10. Data shows mean values ± SE. Significance is denoted by letters.

SGRs for biomass were tested at 40 rather than 100 µmol photons m^-2^ s^-1^ due to spore release at the higher light intensity. The SGRs showed significant differences across nutrient sources (Welch’s p = 0.03; Fig. 5B; Table S6). However, we could only detect a significant difference between thalli grown in F/2 in comparison to VSE (Games-Howell p = 0.02; Table S6B). Despite this, qualitative observations revealed distinct coloration variations based on the nutrient source. Thalli cultivated in VSE appeared pale, while those grown in JS and F/2 media exhibited a dark red color (Fig. 5B).

### Palmaria hecatensis

Growth was observed across all tested temperatures, though the ability to sustain positive SGRs depended on the photoperiod (p < 0.05; Table S7). At 16L:8D, thalli exhibited the highest growth at 4 and 8°C (above 9% daily), significantly surpassing growth at 12 °C, with approximately 3.9% daily (Tukey’s HSD p < 0.05; Fig. 6A, B, and S3; Table S8). Under a 12L:12D photoperiod, 8°C yielded nearly 4% daily SGRs, higher than the approximate 1% observed at 4 and 12°C (Tukey’s HSD p < 0.05; Fig. 6A, C, and S3; Table S7 - S10). In contrast, the 8L:16D photoperiod supported minimal growth, particularly at 4 and 8°C, with SGRs consistently below 1% daily (Fig. 6A, B, and S3). Thalli cultivated at 12°C exhibited contamination more frequently than in other temperature treatments, irrespective of photoperiod.

**Fig. 6.**
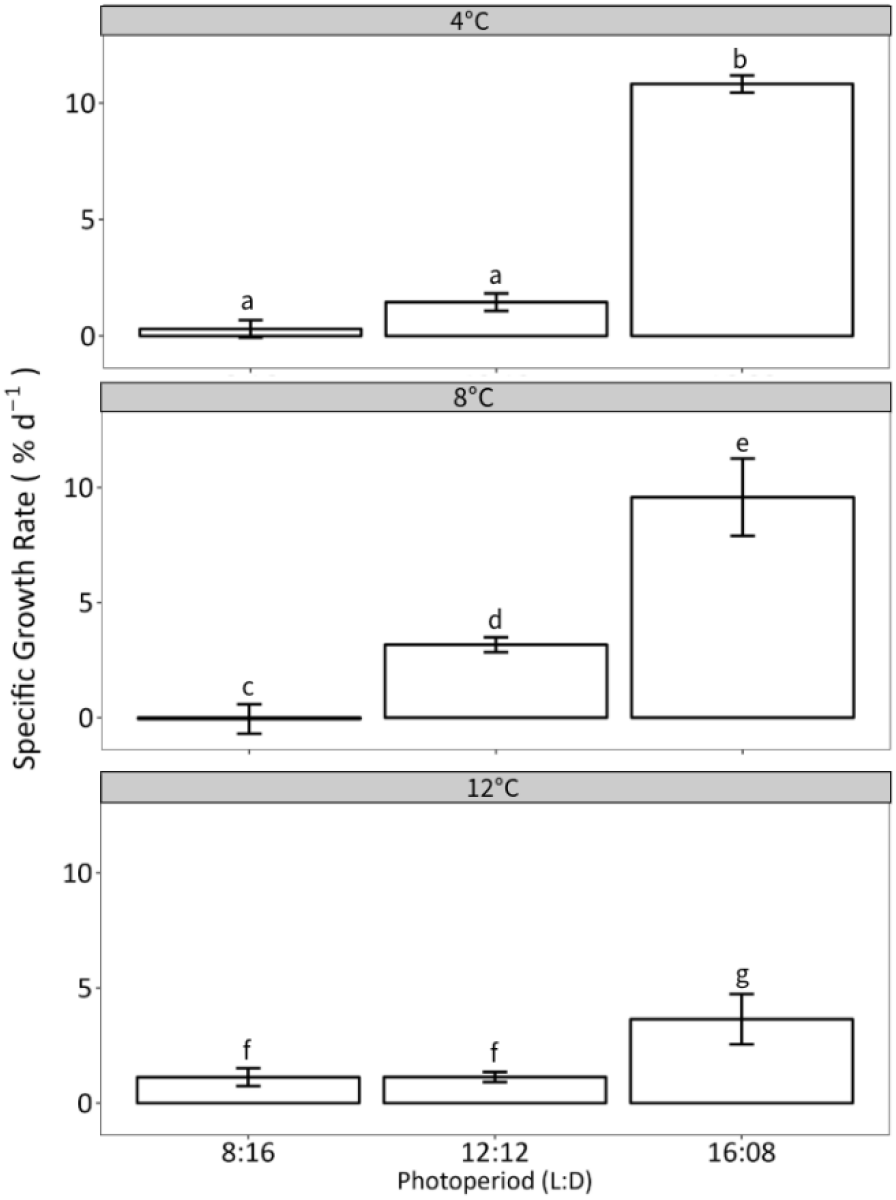
Specific growth rate of *Palmaria hecatensis* as a function of temperature (4, 8, and 12°C) and photoperiod (16L:08D, 12L:12D, and 8L:16D) grown with 40 µmol photons m^-2^ s^-1^ and F/2 as the nutrient source. Data shows mean values ± SE, n = 5. Significance is denoted by letters.

As with *D. mollis, P. hecatensis* exhibited significant differences in SGRs across irradiance levels (p < 0.001; Fig. 7A; Table S11). Thalli exposed to an irradiance of 100 µmol photons m^−2^ s^−1^ achieved the highest SGRs (Tukey’s HSD, p < 0.001; Table S11), followed by those at 40 µmol photons m^−2^ s^−1^, which also demonstrated significantly higher growth than at 20 µmol photons m^−2^ s^−1^ (Tukey’s HSD p < 0.001; Table S11). Unlike *D. mollis, P. hecatensis* showed no fertility or spore release at any irradiance level tested.

**Fig. 7.**
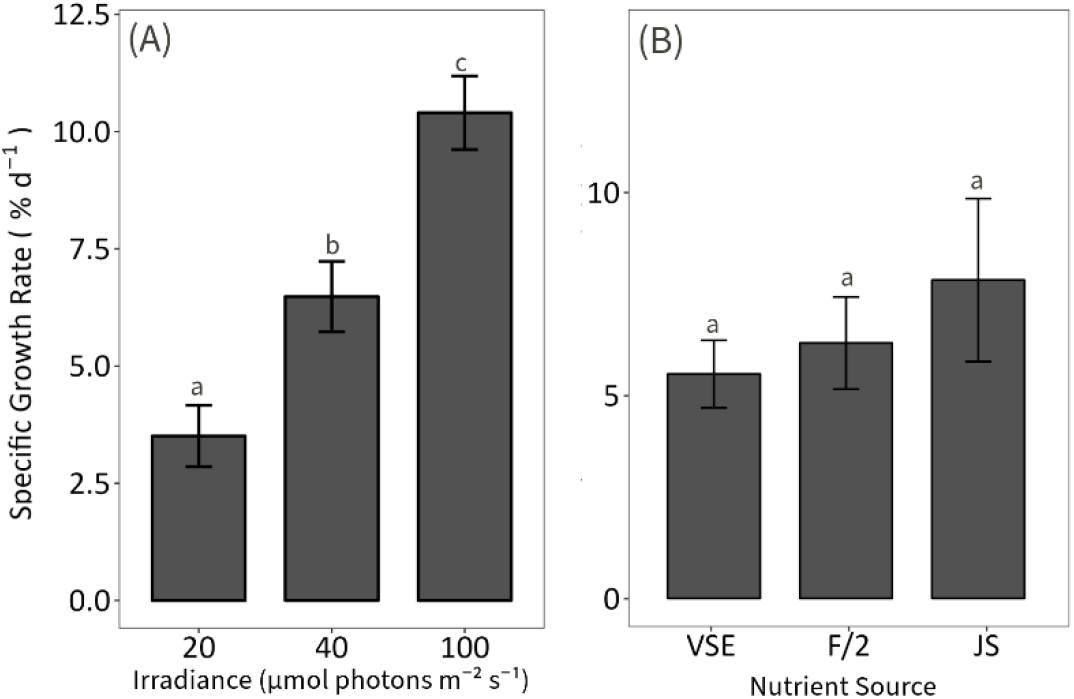
Specific growth rate of *Palmaria hecatensis* as a function of (A) irradiance (20, 40, and 100 µmol photons m^-2^ s^-1^) and (B) nutrient sources (VSE, F/2 and JS), n = 10. Cultures grown at 8°C, 16L:8D, and 40 µmol photons m^-2^ s^-1^. Data shows mean values ± SE. Significance is denoted by letters.

No significant differences in SGRs were observed based on nutrient sources (Fig. 7B; Table S12). Unlike *D. mollis*, the coloration of *P. hecatensis* thalli did not vary among treatments, with all samples maintaining a dark red color throughout the experiment.

## Discussion

Species diversification is essential to enhance the long-term viability, competitiveness, and sustainability of the seaweed farming industry in the United States. By farming various species besides kelp and employing different practices at sea and on land, the industry can possibly reduce its potential vulnerability to disease (Loureiro et al. 2015), market fluctuations (Bindu et al. 2010), and environmental changes (Redmond et al. 2014). This approach also increases opportunities to mitigate risks, enhance economic stability, and promote environmental sustainability by reducing current pressure on wild stocks used as seed source (Huang et al. 2022).

The distinct responses in SGRs of *D. mollis* and *P. hecatensis* to temperature, photoperiod, and irradiance highlight their unique ecological and physiological adaptations despite life history and habitat similarities. *D. mollis* exhibited higher SGRs at warmer temperatures, suggesting a greater thermal tolerance and even a potential preference for warmer conditions. This characteristic likely contributes to its extensive geographical distribution, stretching from Southern California to the Aleutian Islands in Alaska. A range where seasonal water temperatures can vary widely, from 6 to 23°C (Kelley et al. 2011). This thermal flexibility enables *D. mollis* to thrive across diverse marine environments, potentially adapting to the broad temperature fluctuations characteristic of this extensive distribution range.

On the other hand, *P. hecatensis* does not show the same level of increased growth at higher temperatures, suggesting it may have a different thermal optimum to be considered when optimizing the protocols presented here. *P. hecatensis* seems to favor cooler environments or may have a narrower thermal tolerance window preventing its dispersion further south. These results provide insights into the potential effects that water sea surface temperatures may have on the distribution and performance of both species. Moreover, higher irradiance levels triggered spore release in *D. mollis* but not in *P. hecatensis* at the combinations tested. This may indicate a different reproductive strategy or sensitivity to light. The lack of spore release in *P. hecatensis* suggests that it may have a different trigger for reproduction or require even higher light levels than *D. mollis*.

The overall SGRs reported for *D. mollis* here are consistent with those described for *D. mollis* produced in Hawaii and Oregon utilizing flow-through and tumbling setups (Demetropoulos and Langdon 2004a; Demetropoulos and Langdon 2004b). In those studies, growth was observed between 12°C and 17°C, with SGRs peaking at 14°C and achieving an average daily increase of 10% (± 0.5). These results highlight the striking difference between the suitable temperature ranges of Alaskan *D. mollis* and those from lower latitudes. While Alaskan *D. mollis* still grew at 12°C during these experiments, filamentous contaminants frequently proliferated. It is presumed that these contaminants were introduced to the system as endophytes and, therefore, are unlikely to be avoided.

Interestingly, a 12L:12D photoperiod regime is recommended for culture storage for *D. mollis* (Evans 2000). However, our results support that this photoperiod can be used for biomass production, as SGRs were similar between 16L:8D and 12L:12D. This outcome contrasts with other studies on *P. palmata* and *D. mollis*, showing a long-day photoperiod promoting the most growth (Demetropoulos and Langdon 2004b; Pang et al. 2004). A neutral photoperiod at any of the temperatures tested may be used to store cultures of *P. hecatensis*.

*D. mollis* exhibited a daily SGR of 12% ± 1 on average when exposed to 100 µmol photons m^-2^ s^-1^, combined with a temperature of 8°C and a photoperiod of 16L:8D. Such SGRs are consistent with studies conducted on ecotypes from lower latitudes (Demetropoulos and Langdon 2004a; Demetropoulos and Langdon 2004b; Parjikolaei et al. 2013). However, continuous exposure to such irradiance may be unsuitable for cultures with relatively low initial biomass, as spore releases followed by tissue degradation may decrease the efficiency of biomass production. While spore release is well-documented for *P. palmata*, it is relatively understudied for *D. mollis*. It is likely that higher stocking densities of 3-4 kg per m^-2^, as described for on-land cultures in Hawaii, could mitigate the impact of high irradiance on the reproductive cycle of *D. mollis* (Demetropoulos and Langdon 2004b). Until that is tested, and full factorial experiments are completed for the three factors tested here, 40 µmol photons m^-2^ s^-1^ seem to be the most suitable irradiance to promote continuous growth while supporting healthy biomass. An alternative to using lower irradiance levels could be utilizing opaque tanks with high light levels only permeating the surface of the tank (Demetropoulos and Langdon 2004b).

In contrast with *D. mollis, P. hecatensis* showed slower growth when exposed to an 8L:16D regime, particularly at lower temperatures. With only 8 hours of light, *P. hecatensis* individuals may not have enough energy to sustain growth throughout the 24-hour cycle, which could lead to an energy deficit. At lower temperatures, metabolic rates typically decrease, reducing the efficiency of photosynthesis and other physiological processes (Harrison and Hurd 2001). This means that to compensate for lower metabolic rates, *P. hecatensis* would require more light to achieve the same growth as it would at higher temperatures. Thalli of *P. hecatensis* have a leathery texture that resembles that of kelp, contrasting the leafier texture of *D. mollis* (per.obs.). The relatively lower SGR observed on the short-day photoperiod compared to *D. mollis* may be linked to the thickness of the thalli requiring more energy to grow (Hawke and Scagel 1986). However, to our knowledge, this aspect has not been explored.

Previous studies on *D. mollis* co-cultured with abalone showed that supplementation with nitrate promotes more growth in the long term (i.e., nine weeks). In comparison, ammonium is more effective in 2-5 weeks (Demetropoulos and Langdon 2004b). Such results partially align with our finding where the addition of VSE (with nitrate as the source of nitrogen) showed suboptimal biomass performance compared to those supplemented with F/2 (also with nitrate as a source of nitrogen) and JS (with ammonium and nitrate as a source of nitrogen). Up-taking and utilizing nitrate is costly, as energy is needed to assimilate nitrate via ammonium (Jin et al. 1998; Cohen and Fong 2004). The addition of ammonium (NH_4_^+^) likely could contribute to higher protein content by direct incorporation into amino acids, potentially facilitated by ammonium transporters increasing rapid uptake into cells (Lobban and Harrison 1994; Jung et al. *unpublished*).

Furthermore, the observed differences in blade coloration over time show the direct effect of nutrient supplementation on seaweed assimilation and storage capacity. *D. mollis* blades were darker in coloration, potentially from a higher concentration of pigments at the end of the experiment when grown with F/2 and JS. Jung et al. (*unpublished*) experimented further, finding that increased pigment concentration was present when *D. mollis* was exposed to F/2 and JS. Still, this trend did not remain valid for VSE. Seaweed can store excess nitrogen in pigments and other compounds during nutrient-replete events for future growth (McGlathery et al. 1996; Naldi and Wheeler 1999). Jung et al. (*unpublished*) outputs indicate that based on coloration and nitrate alone, the differences observed due to nutrient supplementation cannot be explained. Another contributing factor could be the presence of trace metals such as zinc and copper. Both metals are essential for photosynthesis, cellular metabolism, and growth and, therefore, merit further exploration into their effect on biomass performance (Jung et al. *unpublished*; Demetropoulos and Langdon 2004b; Howarth and Cole 1985).

Finally, *P. hecatensis* might likely have different nutrient uptake mechanisms or efficiencies than *D. mollis* (Harrison and Hurd 2001). This is a topic worth exploring further. Similar to *D. mollis*, phycobiliproteins in thalli exposed to F/2 and JS have shown significantly higher concentrations compared to VSE after 14 days of exposure (Jung et al. *unpublished*). Such outcomes suggest that F/2 and JS formulations likely provide a more enriched nutrient supply, perhaps with a more favorable balance of micro and macronutrients, aside from nitrogen, compared to VSE. This could be responsible for enhancing overall metabolic activity, including synthesizing phycobiliproteins, which are key for light capture and photosynthesis, and hence, growth.

## Conclusions

Results from this study allow for the creation of cultivation protocols that can be refined further to obtain optimal conditions to maximize the growth of *D. mollis* and *P. hecatensis* grown in on-land indoor systems. Although a pairwise approach was used here, optimizing these cultivation protocols would require a full factorial approach. Such an approach would allow the investigation of the interaction effects between the multiple factors tested. In turn, this would provide a deeper understanding of how different variables influence each other and the outcome. It would also provide broader insight into the experimental conditions by simultaneously examining all possible combinations of factors.

Overall, the required adjustments on the environmental parameters to grow *D. mollis* in onland indoor settings and the differences measured between the *D. mollis* and *P. hecatensis* highlight the relevance of tailoring cultivation protocols based on seedstock origin. Our findings also highlight the importance of considering targeted seaweed species’ specific nutritional and storage needs to optimize productivity and quality.

In brief, suitable combinations of temperature, photoperiod, and irradiance to grow *D. mollis* include 8°C with a photoperiod of 16L:8D or 12L:12D. While a photoperiod of 8L: 16D at 4, 8 or 12°C and irradiance of 20 and 40 µmol photons m^-2^ s^-1^ could be used for storage. Further optimization should explore 12L:12D at any of the temperatures tested and 16L:8D at 8°C and 12°C. On the other hand, priority should be placed on 4 and 8°C with a 16L:8D photoperiod to grow *P. hecatensis*, with further protocol optimization comparing these combinations with multiple irradiance levels.

Moreover, further research is essential to understand the nitrogen storage mechanisms and protein compositions of *P. hecatensis*. This understanding could lead to a more efficient use of nutrient sources to increase biomass production. In addition, understanding species-specific responses to nutrient supplementation is also valuable for ecological studies in the context of changing environmental conditions, particularly in rapid warming environments such as those experienced in the Gulf of Alaska, where both species thrive. Finally, the recent range expansion of *P. hecatensis* along the east coast of Russia and Japan may increase the potential for the commercial cultivation of the species (Skriptsova et al. 2023). Therefore, additional research on its ecology and reproductive phenology could greatly improve cultivation practices.

## Declarations

### Availability of Data and Materials

The datasets generated and analyzed during the current study are available from the corresponding author upon reasonable request.

### Code Availability

Not Applicable.

### Author Contributions

**Dittrich:** Conceptualization, Methodology, Formal Analysis, Investigation, Data Curation, Writing – Original Draft, Review and Editing. **Meyer:** Investigation and Writing – Review and Editing. **Stekoll:** Conceptualization, Methodology, Writing – Review and Editing and Supervision. **Kelley:** Methodology, Validation, Writing – Review and Editing and Supervision. **Umanzor:** Conceptualization, Methodology, Resources, Writing – Original Draft, Writing – Review and Editing, Supervision and Funding Acquisition.

## Acknowledgements

The authors would like to acknowledge that this work was produced on the ancestral lands of the Tlingit people to whom we are grateful.

