## Supplementary Materials for "Cultivation protocols for the rhodophytes, *Devaleraea mollis* and *Palmaria hecatensis* from Alaska"

### Supplementary Figures and Tables

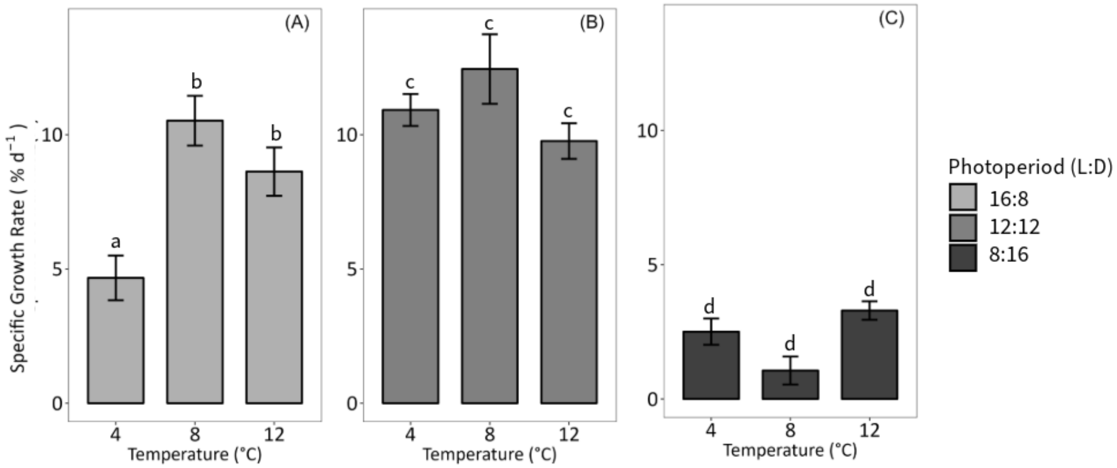


**Fig. S1** Specific growth rate of *Devaleraea mollis* as a function of photoperiod (16L:08D, 12L:12D, and 8L:16D) and temperature (4, 8, and 12°C) grown with and 40 μmol photons m$^{-2}$ s$^{-1}$ and F/2 as the nutrient source. Data shows mean values ± SE, n = 5. Significance is denoted by letters.

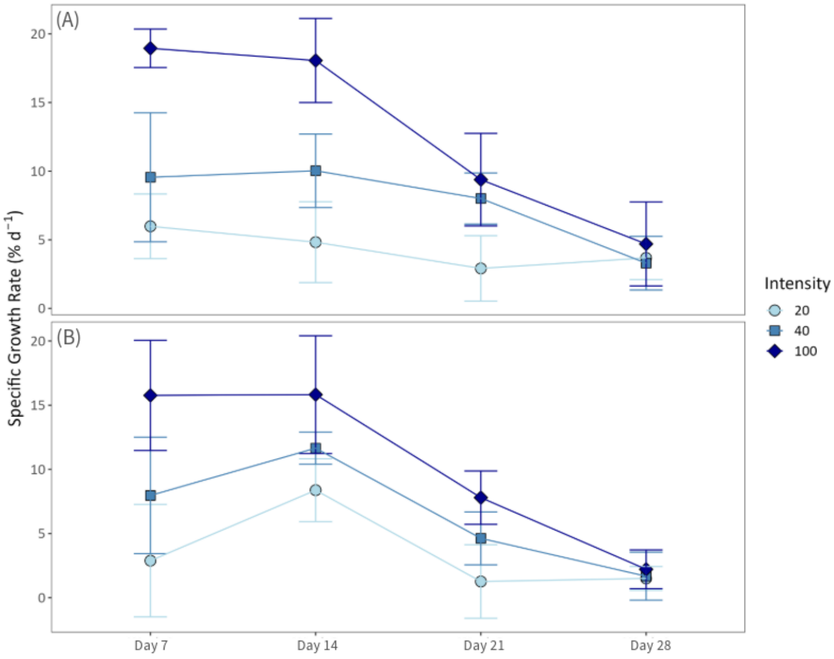


**Fig. S2** Weekly SGRs as a function of irradiance in μmol photons m$^{-2}$ s$^{-1}$ for (A) *Devaleraea mollis* and (B) *Palmaria hecatensis*. Cultures were grown at 8°C with a 16L:08D, and F/2 for nutrients. Data shows mean values ± SD, n = 5.

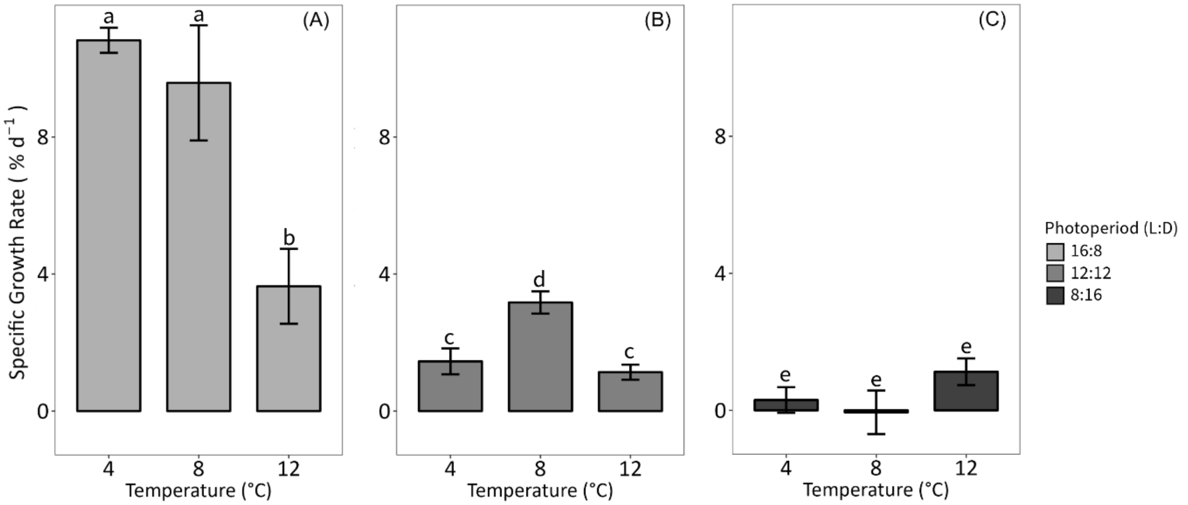


**Fig. S3** Specific growth rate of *Palmaria hecatensis* as a function of photoperiod (16L:08D, 12L:12D, and 8L:16D) and temperature (4, 8, and 12°C) grown with and 40 μmol photons $m^{-2}$ $s^{-1}$ and F/2 as the nutrient source. Data shows mean values ± SE, n = 5. Significance is denoted by letters.

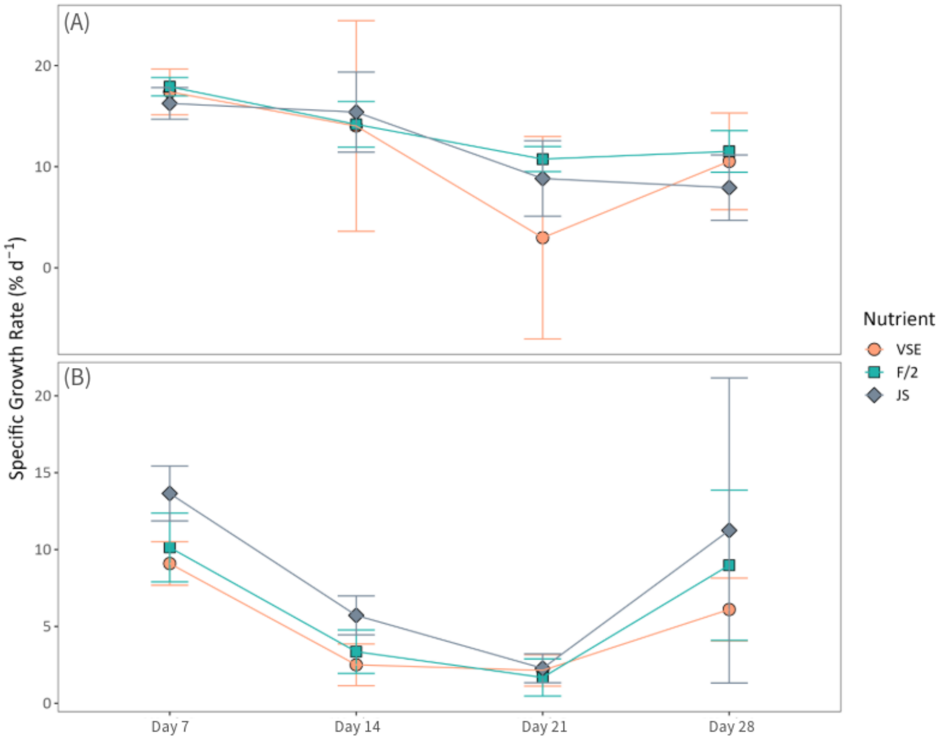


**Fig. S4** Weekly SGRs as a function of nutrient source for (A) *Devaleraea mollis* and (B) *Palmaria hecatensis*. Cultures were grown at 8°C, 16L:08D, and 40 μmol photons $m^{-2}$ $s^{-1}$ for *Devaleraea mollis* and 100 μmol photons $m^{-2}$ $s^{-1}$ for *Palmaria hecatensis*. Data shows mean values ± SD, n = 5.

### Supplementary Tables

**Table S1.** One-way ANOVA for the effect photoperiod (8L:16D, 12L:12D, and 16L:8D) on specific growth rates of *Devaleraea mollis* at three different temperatures (4, 8, and 12°C). Asterisks indicate a significant effect.

| Temperature: 4°C | | | | | |
|---|---|---|---|---|---|
| **Factor** | **df** | **Sum Sq** | **Mean Sq** | **F value** | **p-value** |
| Photoperiod | 2 | 190.97 | 95.48 | 44.56 | 2.79e-06* |
| Residuals | 12 | 25.71 | 2.14 | | |
| **Temperature: 8°C** | | | | | |
| **Factor** | **df** | **Sum Sq** | **Mean Sq** | **F value** | **p-value** |
| Photoperiod | 2 | 371.3 | 185.67 | 39.71 | 5.11e-06* |
| Residuals | 12 | 56.1 | 4.68 | | |
| **Temperature: 12°C** | | | | | |
| **Factor** | **df** | **Sum Sq** | **Mean Sq** | **F value** | **p-value** |
| Photoperiod | 2 | 119.19 | 59.59 | 26.12 | 4.25e-05* |
| Residuals | 12 | 27.38 | 2.28 | | |

**Table S2.** Tukey's HSD examining differences in the specific growth rates of *Devaleraea mollis* as a function of photoperiod (8L:16D, 12L:12D, and 16L:8D) at three different temperatures (4, 8, and 12°C). Asterisks indicate a significant effect.

| Temperature: 4°C | | | | |
|---|---|---|---|---|
| **Comparison** | **Difference** | **Lower CI (lwr)** | **Upper CI (upr)** | **Adjusted p-value (p adj)** |
| 16:08 - 12:12 | -6.25513 | -8.724965 | -3.785296 | 0.0000558* |
| 8:16 - 12:12 | -8.413918 | -10.883752 | -5.944084 | 0.0000028* |
| 8:16 - 16:08 | -2.158787 | -4.628621 | 0.311047 | 0.0894072 |
| **Temperature: 8°C** | | | | |
| **Comparison** | **Difference** | **Lower CI (lwr)** | **Upper CI (upr)** | **Adjusted p-value (p adj)** |
| 16:08 - 12:12 | -1.922669 | -5.571009 | 1.72567 | 0.3687246 |
| 8:16 - 12:12 | -11.383925 | -15.032265 | -7.735586 | 0.0000069* |
| 8:16 - 16:08 | -9.461256 | -13.109596 | -5.812916 | 0.0000443* |
| **Temperature: 12°C** | | | | |
| **Comparison** | **Difference** | **Lower CI (lwr)** | **Upper CI (upr)** | **Adjusted p-value (p adj)** |
| 16:08 - 12:12 | -1.138911 | -3.687596 | 1.409774 | 0.4798067 |
| 8:16 - 12:12 | -6.467232 | -9.015917 | -3.918547 | 0.0000547* |
| 8:16 - 16:08 | -5.328321 | -7.877006 | -2.779636 | 0.0003276* |

**Table S3.** One-way ANOVA for the effect photoperiod (8L:16D, 12L:12D, and 16L:8D) on specific growth rates of *Devaleraea mollis* at three different temperatures (4, 8, and 12°C). Asterisks indicate a significant effect.

| Photoperiod: 8L:16D | | | | | |
|---|---|---|---|---|---|
| **Factor** | **df** | **Sum Sq** | **Mean Sq** | **F value** | **p-value** |
| Temperature | 2 | 12.86 | 6.428 | 6.149 | 0.0145* |
| Residuals | 12 | 12.54 | 1.045 | | |
| Photoperiod: 12L:12D | | | | | |
| **Factor** | **df** | **Sum Sq** | **Mean Sq** | **F value** | **p-value** |
| Temperature | 2 | 18.08 | 9.042 | 2.193 | 0.154 |
| Residuals | 12 | 49.47 | 4.123 | | |
| Photoperiod: 16L:8D | | | | | |
| **Factor** | **df** | **Sum Sq** | **Mean Sq** | **F value** | **p-value** |
| Temperature | 2 | 89.22 | 44.61 | 11.35 | 0.00171* |
| Residuals | 12 | 47.18 | 3.93 | | |

**Table S4.** Tukey's HSD examining differences in the specific growth rates of *Devaleraea mollis* as a function of photoperiod (8L:16D, 12L:12D, and 16L:8D) at three different temperatures (4, 8, and 12°C). Asterisks indicate a significant effect.

| Photoperiod: 8L:16D | | | | |
|---|---|---|---|---|
| **Comparison** | **Difference** | **Lower CI (lwr)** | **Upper CI (upr)** | **Adjusted p-value (p adj)** |
| 8 - 4 | -1.4479802 | -3.1731693 | 0.2772089 | 0.1045146 |
| 12 - 4 | 0.7873909 | -0.9377982 | 2.51258 | 0.4657695 |
| 12 - 8 | 2.2353711 | 0.510182 | 3.9605602 | 0.0122077* |
| Photoperiod: 12L:12D | | | | |
| **Comparison** | **Difference** | **Lower CI (lwr)** | **Upper CI (upr)** | **Adjusted p-value (p adj)** |
| 8 - 4 | 1.522027 | -1.903974 | 4.9480294 | 0.4836696 |
| 12 - 4 | -1.159295 | -4.585297 | 2.2667071 | 0.6488754 |
| 12 - 8 | -2.681322 | -6.107324 | 0.7446797 | 0.1341397 |
| Photoperiod: 16L:8D | | | | |
| **Comparison** | **Difference** | **Lower CI (lwr)** | **Upper CI (upr)** | **Adjusted p-value (p adj)** |
| 8 - 4 | 5.854489 | 2.5089703 | 9.200007 | 0.0014541* |
| 12 - 4 | 3.956924 | 0.6114061 | 7.302443 | 0.0209804* |
| 12 - 8 | -1.897564 | -5.2430826 | 1.447954 | 0.3194977 |

**Table S5 (a) & (b):** One-way ANOVA for irradiance experiments for *Devaleraea mollis* (a), with a pairwise comparison post hoc analysis (b). Asterisks indicate a significant effect.

| (a) | | | | | |
|---|---|---|---|---|---|
| **Factor** | **df** | **Sum Sq** | **Mean Sq** | **F value** | **p-value** |
| Irradiance | 2 | 179.64 | 89.82 | 35.54 | <0.001* |
| Residuals | 12 | 30.33 | 2.53 | | |

| (b) | | | | | |
|---|---|---|---|---|---|
| **Factor** | **Estimate** | **SE** | **df** | **t.ratio** | **p-value** |
| 20 - 40 irradiance | -3.37 | 1.01 | 12 | -3.53 | 0.0147* |
| 20 - 100 irradiance | -8.42 | 1.01 | 12 | -8.37 | <0.0001* |
| 40 - 100 irradiance | -5.05 | 1.01 | 12 | -5.02 | 0.0008* |

**Table S6 (a) & (b):** Welch's One-way ANOVA for the nutrient source experiments for *Devaleraea mollis* (a), with a pairwise t-test with Bonferroni correction (b). Asterisks indicate a significant effect.

| (a) | | | |
|---|---|---|---|
| **Factor** | **df** | **F value** | **p-value** |
| Nutrient Source | 2 | 5.72 | 0.039* |

| (b) | | | |
|---|---|---|---|
| **Group1** | **Group2** | **Mean_Diff** | **p-value** |
| VSE | F/2 | 2.35785 | 0.0204* |
| VSE | JS | 0.87171 | 0.4961 |
| F/2 | JS | 1.48614 | 0.2204 |

**Table S7.** One-way ANOVA for the effect of temperature (4, 8, and 12°C) on specific growth rates of *Palmaria hecatensis* at three different photoperiods (8L:16D, 12L:12D, and 16L:8D). Asterisks indicate a significant effect.

| Photoperiod: 8L:16D | | | | | |
|---|---|---|---|---|---|
| **Factor** | **df** | **Sum Sq** | **Mean Sq** | **F value** | **p-value** |
| Temperature | 2 | 3.677 | 1.838 | 1.585 | 0.245 |
| Residuals | 12 | 13.917 | 1.16 | | |
| Photoperiod: 12L:12D | | | | | |
| **Factor** | **df** | **Sum Sq** | **Mean Sq** | **F value** | **p-value** |
| Temperature | 2 | 11.987 | 5.993 | 12.13 | 0.00132* |
| Residuals | 12 | 5.931 | 0.494 | | |
| Photoperiod: 16L:8D | | | | | |
| **Factor** | **df** | **Sum Sq** | **Mean Sq** | **F value** | **p-value** |
| Temperature | 2 | 147.25 | 73.63 | 10.62 | 0.00222* |
| Residuals | 12 | 83.21 | 6.93 | | |

**Table S8.** Tukey's HSD examining differences on the specific growth rates of *Palmaria hecatensis* as a function of temperature (4, 8, and 12°C) at three different photoperiods (8L:16D, 12L:12D, and 16L:8D). Asterisks indicate a significant effect.

| Temperature: 4°C | | | | |
|---|---|---|---|---|
| **Comparison** | **Difference** | **Lower CI (lwr)** | **Upper CI (upr)** | **Adjusted p-value (p adj)** |
| 16:08 - 12:12 | 9.372095 | 7.964074 | 10.7801154 | <0.0001* |
| 8:16 - 12:12 | -1.148417 | -2.556438 | 0.2596038 | 0.1161019 |
| 8:16 - 16:08 | -10.520512 | -11.928532 | -9.1124908 | <0.0001* |
| Temperature: 8°C | | | | |
| **Comparison** | **Difference** | **Lower CI (lwr)** | **Upper CI (upr)** | **Adjusted p-value (p adj)** |
| 16:08 - 12:12 | 6.408864 | 2.427966 | 10.3897619 | 0.0027633* |
| 8:16 - 12:12 | -3.225623 | -7.206521 | 0.7552747 | 0.1188724 |
| 8:16 - 16:08 | -9.634487 | -13.615385 | -5.6535892 | 0.000086* |
| Temperature: 12°C | | | | |
| **Comparison** | **Difference** | **Lower CI (lwr)** | **Upper CI (upr)** | **Adjusted p-value (p adj)** |
| 16:08 - 12:12 | 2.506834247 | -0.06621531 | 5.0798838 | 0.0563951 |
| 8:16 - 12:12 | -0.008810131 | -2.58185969 | 2.56423943 | 0.999954 |
| 8:16 - 16:08 | -2.515644378 | -5.08869394 | 0.05740518 | 0.0555014 |

**Table S9.** One-way ANOVA for the effect photoperiod (8L:16D, 12L:12D, and 16L:8D) on specific growth rates of *Palmaria hecatensis* at three different temperatures (4, 8, and 12°C). Asterisks indicate a significant effect.

| Photoperiod: 8L:16D | | | | |
|---|---|---|---|---|
| **Comparison** | **Difference** | **Lower CI (lwr)** | **Upper CI (upr)** | **Adjusted p-value (p adj)** |
| 8 - 4 | -0.3584182 | -2.1755284 | 1.458692 | 0.8601384 |
| 12 - 4 | 0.8241112 | -0.99299 | 2.641221 | 0.4699841 |
| 12 - 8 | 1.1825294 | -0.6345808 | 2.99964 | 0.2323067 |
| Photoperiod: 12L:12D | | | | |
| **Comparison** | **Difference** | **Lower CI (lwr)** | **Upper CI (upr)** | **Adjusted p-value (p adj)** |
| 8 - 4 | 1.7187882 | 0.532608 | 2.9049685 | 0.0058777* |
| 12 - 4 | -0.3154957 | -1.501676 | 0.8706845 | 0.7625709 |
| 12 - 8 | -2.0342839 | -3.220464 | -0.8481037 | 0.0017044* |
| Photoperiod: 16L:8D | | | | |
| **Comparison** | **Difference** | **Lower CI (lwr)** | **Upper CI (upr)** | **Adjusted p-value (p adj)** |
| 8 - 4 | -1.244443 | -5.687611 | 3.198726 | 0.7409608 |
| 12 - 4 | -7.180756 | -11.62394 | -2.737588 | 0.0026846* |
| 12 - 8 | -5.936314 | -10.379482 | -1.493145 | 0.0100648* |

**Table S10.** Tukey's HSD examining differences in the specific growth rates of *Palmaria hecatensis* as a function of photoperiod (8L:16D, 12L:12D, and 16L:8D) at three different temperatures (4, 8, and 12°C). Asterisks indicate a significant effect.

| Temperature: 4°C | | | | | |
|---|---|---|---|---|---|
| **Factor** | **df** | **Sum Sq** | **Mean Sq** | **F value** | **p-value** |
| Photoperiod | 2 | 333.1 | 166.5 | 239.1 | 2.15e-10* |
| Residuals | 12 | 8.4 | 0.7 | | |
| Temperature: 8°C | | | | | |
| **Factor** | **df** | **Sum Sq** | **Mean Sq** | **F value** | **p-value** |
| Photoperiod | 2 | 240.5 | 120.25 | 21.6 | 0.000105* |
| Residuals | 12 | 66.8 | 5.57 | | |
| Temperature: 12°C | | | | | |
| **Factor** | **df** | **Sum Sq** | **Mean Sq** | **F value** | **p-value** |
| Photoperiod | 2 | 21.02 | 10.511 | 4.52 | 0.0344* |
| Residuals | 12 | 27.91 | 2.325 | | |

**Table S11 (a) & (b):** One-way ANOVA for irradiance experiments for *Palmaria hecatensis* (a), with a pairwise comparison post hoc analysis (b). Asterisks indicate a significant effect.

| (a) | | | | | |
|---|---|---|---|---|---|
| **Factor** | **df** | **Sum Sq** | **Mean Sq** | **F value** | **p-value** |
| Irradiance | 2 | 179.64 | 89.82 | 35.54 | <0.001* |
| Residuals | 12 | 30.33 | 2.53 | | |

| (b) | | | | | |
|---|---|---|---|---|---|
| **Factor** | **Estimate** | **SE** | **df** | **t.ratio** | **p-value** |
| 20 - 40 irradiance | -2.97 | 1.01 | 12 | -2.87 | 0.0349* |
| 20 - 100 irradiance | -6.89 | 1.01 | 12 | -6.65 | <0.0001* |
| 40 - 100 irradiance | -3.92 | 1.01 | 12 | -3.78 | 0.0068* |

**Table S12 (a) & (b):** Welch's One-way ANOVA for the nutrient source experiments for *Palmaria hecatensis* (a), with a pairwise t-test with Bonferroni correction (b). Asterisks indicate a significant effect.

| (a) | | | | | |
|---|---|---|---|---|---|
| **Factor** | **df** | **Sum Sq** | **Mean Sq** | **F value** | **p-value** |
| Irradiance | 2 | 179.64 | 89.82 | 35.54 | <0.001* |
| Residuals | 12 | 30.33 | 2.53 | | |

| (b) | | | | | |
|---|---|---|---|---|---|
| **Factor** | **Estimate** | **SE** | **df** | **t.ratio** | **p-value** |
| 20 - 40 irradiance | -2.97 | 1.01 | 12 | -2.87 | 0.0349* |
| 20 - 100 irradiance | -6.89 | 1.01 | 12 | -6.65 | <0.0001* |
| 40 - 100 irradiance | -3.92 | 1.01 | 12 | -3.78 | 0.0068* |
